# Immune escape facilitation by mutations of epitope residues in RdRp of SARS-CoV-2

**DOI:** 10.1101/2021.11.18.469065

**Authors:** Aayatti Mallick Gupta, SasthiCharan Mandal, Sukhendu Mandal, Jaydeb Chakrabarti

## Abstract

SARS-CoV-2 has considerably higher mutation rate. SARS-CoV-2 possesses a RNA dependent RNA polymerase (RdRp) which helps to replicate its genome. The mutation P323L in RdRp is associated with the loss of a particular epitope (321-327) from this protein which may influence the pathogenesis of the concern SARS-CoV-2 through the development of antibody escape variants. We consider the effect of mutations in some of the epitope regions including the naturally occurring mutation P323L on the structure of the epitope and their interface with paratope using all-atom molecular dynamics (MD) simulation studies. P323L mutations cause conformational changes in the epitope region by opening up the region associated with increase in the radius of gyration and intramolecular hydrogen bonds, making the region less accessible. Moreover, the fluctuations in the dihedral angles in the epitope:paratope (IgG) interface increase which destabilize the interface. Such mutations may help in escaping antibody mediated immunity of the host.

## 1. Introduction

RNA viruses are prone to mutations, depending on the fidelity of the RNA polymerase. These mutations direct the overall variations in the viral genome in the population [1,2] and are beneficial for viral adaptation [3,4]. SARS-CoV-2 is an enveloped, single and positive stranded RNA virus belonging to the genus beta corona virus [5]. This genus includes two more RNA viruses, like SARS-CoV [6] and MERS-CoV [7], responsible for epidemics, like the Severe Acute Respiratory Syndrome (SARS) and the Middle East Respiratory Syndrome (MERS), respectively. As a common characteristic of RNA viruses, the more the SARS-CoV-2 is spreading across the globe, newer mutational hotspots are emerging in its genome. Lack of pre-existing immunity against a particular variant and its high transmissibility leave us vulnerable to infection that leads the WHO to the announcement of a worldwide public health emergency [8,9]. The microscopic mechanism of immune escape by the mutated variants remains elusive. The emerging SARS-CoV-2 mutant variants include RNA dependent RNA polymerases (RdRp) as one of the mutation hot spots [10]. RdRp is a multi-domain protein that forms key transcription machinery in cooperation with cofactors nsp7 and nsp8 to carry out the proof-reading activity [11-13]. Recurrent P323L mutation of RdRp across the world has been reported [14]. Cryo EM structural study obtained for SARS-CoV-2 RdRp [15] has tempted to speculate the conformation changes due to its mutation. In an earlier work [16] we show that the region 321-327 in RdRp forms an epitope of the antigen that is recognizable by the paratope of the antibody of the human host cell, while P323L mutation is associated with loss in epitope. Such loss of antigenic determinant impairing immune recognition has been found in many viral disease models [17]. So, the question is: Does the loss in epitope in RdRp suffice to cause impairment of the paratope interaction? We address this question in the present work.

The relevant antibody for the epitope region 321-327 is not known. A high concentration of IgG antibody has been observed from week 3 to week 7 after SARS-CoV-2 viral exposure indicating the immune response against the virus [18]. Antigen specific antibody IgM could not be detectable after illness onset suggesting the possibility that most of the antigen specific antibody class switching to IgG occurs within 1 week after infection [19]. The effects of RdRp mutations on the binding of the epitope to IgG paratope is of paramount interest. The immune response involves several mechanisms of molecular recognition and interaction between epitope and paratope. Biomolecular binding typically involves changes in conformation which is associated with free energy and entropy costs [20]. The thermodynamic costs between two conformational (complex versus free states) states can be estimated by comparing the marginal dihedral distributions in the two conformations. The ratio of the peaks in the distributions gives population difference associated with the Boltzmann factor of the underlying free energy, while the entropy is given using the Gibb’ s formula [20]. All the dihedral contributions can be added to get the changes in a given residue and overall the residues for the total changes. If the correlations between the binding partners are ignored, one can speak in terms of changes in the thermodynamic quantities of individual components. A negative change in free energy and entropy represent conformational stability and ordering in a given state with respect to the reference state. We base our analysis on changes in conformational thermodynamics, if the changes enhance the stability and order in the epitope-paratope interface.

We perform all-atom molecular dynamics (MD) simulations on the wild type (wt) and mutated type (mt) protein in their free and antibody bound states using theGROMOS96 53a6 force-field in the GROMACS 2018.6 package. We study the structural changes associated with the mutation and compute the corresponding thermodynamic changes. This is done by comparing the dihedral distributions in the mutated RdRp conformation for the wild type RdRp conformation. We further consider both mutated and wild type RdRp in presence of IgG antibody to study the relative conformational stability and order the complexes. Our observations are as follows: (1) We consider a few mutations, namely, the epitope residues P323L, F324L and T326L. We observe that the epitope region undergoes conformational disorder due to the mutations. (2) The wild type RdRp shares a conformationally stable interface with the antibody, whereas mutated RdRp does not. Thus, the mutated RdRp cannot form a stable interface with IgG compared to the wild type protein. Antigen binding to the antibody is vital in immune response and has several biomedical applications including vaccines and immune-therapeutics [21,22].

## 2. Materials and methods

### 2.1 System preparation

We have considered the cryo-EM structure of SARS-CoV-2 RNA-dependent RNA polymerase (PDB id: 6M71, chain A) [15] to study the effect of mutation on structural stability. The mutations are obtained from the cryo-EM structure. The epitope is identified using a 5-fold cross-validation approach considering computed volume, hydrophobicity, polarity, together with the relative surface accessibility and secondary structure [23]. IgG antibody (PDB id: 2DD8, chain H and L) [24] is used for paratope prediction using RdRp as antigen with antibody i-patch program [25]. Antibody i-Patch uses antibody-specific statistics to annotate residues with a score indicating how likely they are to be in contact with the supplied antigen. Antibody i-patch Scores range from 0 to 12, with residues higher values corresponding to greater binding likelihood with antigen. Thus, the complementarity determining region (CDR) according to Immunogenetics Information System (IMGT) [26, 27] definition can be located.

### 2.2 Docking studies

Antigen-antibody docking studies have been carried out using the Antibody Mode of the ClusPro server (www.cluspro.bu.edu) [28]. CDR residues of an antibody are defined while the rest of the residues that does not fall in the CDR are masked. We have performed docking separately for IgG antibody with wildtype and mutated RdRp protein respectively. Vina1.1.2 [29] has been utilized to perform molecular docking with the drug molecule and RdRp to predict the effect of the later on drug susceptibility due to P323L mutation. The docking boxes have been set at the hydrophobic cleft close to P323L mutation site. Two separate dockings have been carried out using (i) wild type RdRp and Tegobuvir (drug molecule) and (ii) mutated RdRp and Tegobuvir. The search exhaustiveness is set as 32, and the number of binding modes is set as 9.

### 2.3. MD simulation

We perform molecular dynamics simulations in explicit water with counter-ions using GROMOS96 53a6 force-field [30]. spc216 (simple point charge) water molecule [31] is used for solvation under a cubic box with a minimum distance between the protein and the box 1.0 Å. Chlorine and sodium counter ions are added as required replacing the water molecules to neutralize the positive or negative charge. Particle mesh Ewald (PME) sum method has been used for treating the long-range electrostatic interactions [32]. The energy minimization has been performed until the maximum force of the system became < 100 kJ/mol/nm to relieve steric clashes and inappropriate geometry. For equilibration of the systems, a short NVT equilibration phase with the heavier atoms fixed and then follows a longer NPT runs. The systems are subjected to isothermal (300K) and isobaric (1 atm) conditions under periodic boundary conditions and 2 fs time-step. The total number of particles (N= 254640) including water, pressure and temperature have been fixed for each of the systems to make the simulated ensembles equivalent. The simulation trajectories are calculated up to 500 ns. The equilibrations of the simulated structures are assessed from the root-mean square deviation (RMSD). RMSD determines the structural changes over time. The RMSD data (Supplementary information, SI Figs. S1(A)-(E)) show that the systems are equilibrated beyond 300ns. We analyze the data from the last 200 ns trajectory. Various analyses are carried out with tools in GROMACS to examine the system properties. Conformational thermodynamics changes for (a) P323L mutation in RdRp and (b) epitope-paratope interface are estimated from equilibrium fluctuations of the dihedral angles using the Histogram Based Method (HBM) [20]. The histograms of the dihedrals are computed considering all the structures from the equilibrated trajectory (between 300-500ns). 100 samples each with 1000 conformations are generated from the equilibrated 300-500 ns trajectory for each wildtype and mutated system. The final structure of each of the systems given by a snapshot is also shown as SI Fig. S2.

## 3. Results and discussions

The epitope regions of RdRp include residues F321, P322, P323, T324, S325, F326 and G327 [16]. For the given epitope residues(Figs. 1(a) and (b)), the paratope residues are identified using the standard immuno-informatics approach [see methods for details]. The paratope region of IgG as an antibody includes S31, Y32 and T33 at H1 region and H2 region consist of L55, I57, N59 and Y60. R94, D95, T96, G99, D101 and V102 are typically present in the H3 region(Fig. 1c). In the light chain, K31 and S32 comprise the L1 region, Y49-P55 forms the L2 region and L3 region consist of W91-Y97. Docking using the crystal structures of wild type (wt-) RdRp and IgG, biased by the identified epitope and paratope residues yields the paratope-epitope interface within 0.5 to 1 nm distance in the wt-RdRp-IgG complexes shown in Table 1. The interface predominantly consists of the epitope residues P323, F324 and T326. We consider point mutations, like the reported one P323L, and additional mutations F324L and T326L. We denote the mutated systems by mt(P323L)-RdRp, mt(T324L)-RdRp and mt(F326L)-RdRp respectively. In all the cases, we have replaced the residue with hydrophobic residue L, which has a less preference to be epitope residue [33]. The MD simulation studies are carried out in two stages: First, we consider the changes in the epitope conformation under mutation and thermodynamics quantities associated with these changes. Next, we consider the changes in the epitope-paratope region due to mutation.

**Table1:** The interacting residues at the epitope:paratope interface

| System | Epitope | Paratope residue | Nature of interaction |
| --- | --- | --- | --- |
| wt-RdRp-IgG | T324 | H-Y32 | Polar |
|  | P323 | H-G99 | Van der Waals |
|  | T324 | H-D95 | Polar-charged |
|  | T324 | H-Y96 | Polar |
|  | P323 | L-S32 | Hydrogen |
|  | T324 | L-S32 | Polar |
|  | F326 | L-S32 | Hydrogen |
|  | P323 | L-Y49 | Van der Waals |
|  | P323 | L-D50 | Van der Waals |
|  | P323 | L-W91 | Van der Waals |
|  | T324 | L-W91 | Polar |
|  | T324 | L-Y96 | Polar |
|  | F326 | L-W91 | Van der Waals |
|  | F326 | L-S93 | Van der Waals |
|  | F326 | L-S94 | Van der Waals |
| mt(P323L)-RdRp-IgG | T324 | H-G99 | Van der Waals |
|  | F326 | L-W91 | Van der Waals |

**Figs. 1:**
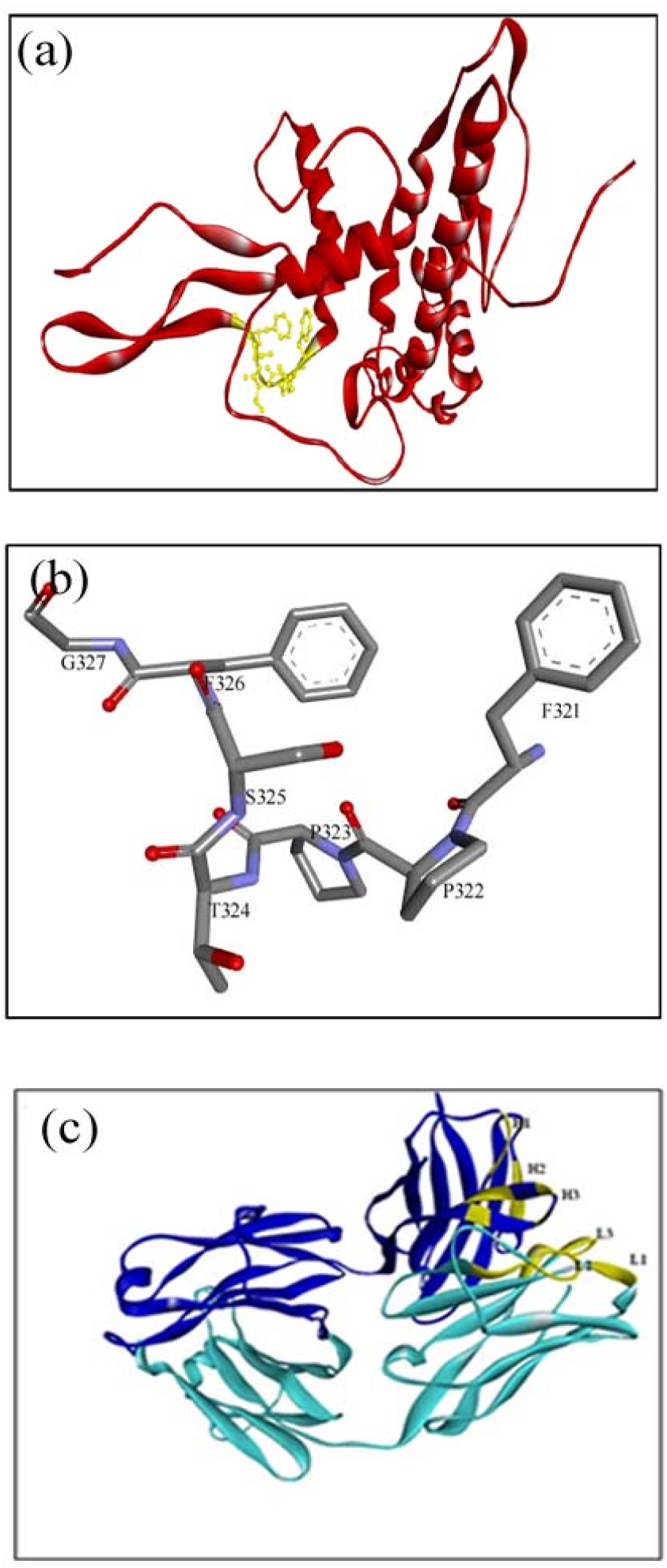
(a) RdRp protein (red) with the epitope region highlighted in yellow.(b) Close up view of the epitope region (ball and stick model) (c) IgG with H chain (dark blue) and L chain (light blue). The paratope regions are shown in yellow.

### 3.1. Conformation changes in epitope due to point mutations

Let us consider in detail the conformations of wt-RdRp and mt(P323L)-RdRp. We characterize epitope conformation using the cosine of the angle *θ* between the vectors joining C_α_atoms of F321 and T324 and G327. We show the distribution of cos*θ*, H(cos*θ*) over the equilibrium trajectories for both the wild type and the mutated proteins in Fig. 2a. The wt-RdRp reveals a sharp peak at *θ*= 90°. In mt(P323L)-RdRp, there is a shift to *θ*= 180°. This represents the opening of the epitope region due to P323L mutation. The changes of conformation are manifested in the other quantities as well. Radius of gyration (R_g_), a measure for the compactness of the structure has been calculated using the standard method as given at Gromacs manual [30]. The distribution H(*R*_*g*_), shown in Fig. 2b, has a peak for larger *R*_*g*_ in mt(P323L)-RdRp than wt-RdRp, depicting lack of compactness associated with loss of epitope region due to P323L mutation. Solvent accessible surface area, S is given by the area of the surface of RdRp with radius *r*_*vdW*_+*r*_*sol*_, where the center of a spherical solvent molecule or probe (*r*_*sol*_ in contact with the atomic van der Waals sphere *r*_*vdW*_. The histogram distribution of solvent accessible area (SASA), H(SASA) (Fig. 2c) shows that the SASA in mt(P323L)-RdRp is less than that in wt-RdRp. Hydrogen bond analysis gives the number of hydrogen bonds at a distance of less than 3.5 Å between all possible donors D and acceptors A with a D-H-A angle of 180° to 30°. The distribution of number of hydrogen bond in the epitope region, H(hb) is higher in mt(P323L)-RdRp than in wt-RdRp (Fig. 2d). We have used the DSSP [34] to assign the secondary structural elements to the residues. Secondary structural changes due to P323L mutation of RdRp from the simulated trajectory reveal the predominance of 5-helix over beta bridge (Figs. 2e and 2f). The change of secondary structural elements in the epitope residue is consistent to epitope loss due to P323L mutation as epitope residues are typically rich in loops and depleted of strands and helixes [33,35].

**Figs. 2:**
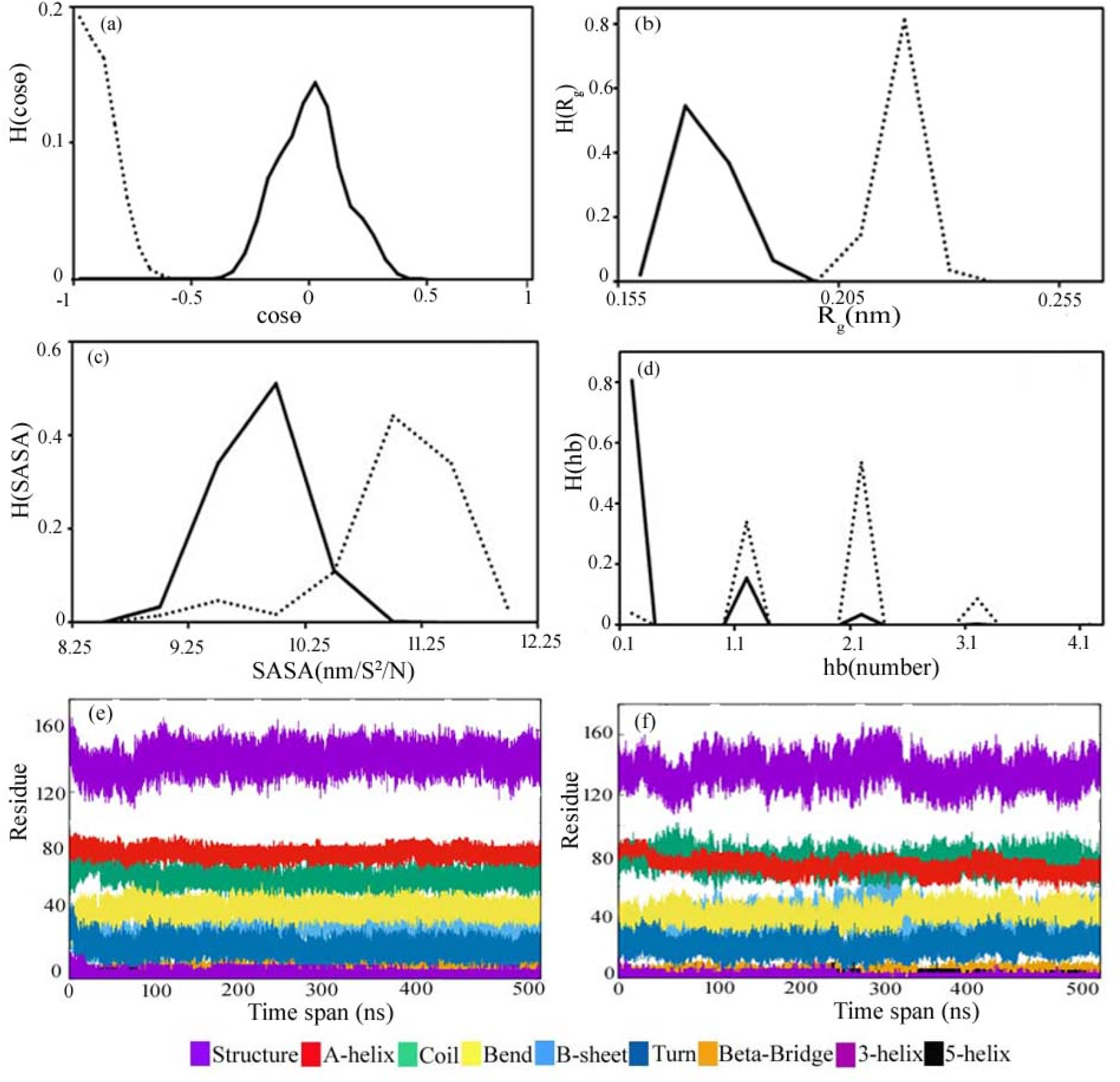
Comparative structural analysis of epitope region in wt-RdRp(solid line) and mt(P323L)-RdRp(dotted line): (a) histogram of cos*θ*, H(cos*θ*) where *θ* is the angle between the vectors joining C_α_atoms of F321 and T324 and G327. (b) Histogram H(R_g_) of the radius of gyration. (c) Histogram H(SASA) of accessible surface area. (d) Histogram H(hb) of the number of hydrogen bond due to intramolecular interactions. Secondary structure with time plot: (e) wt-RdRp and (f) mt(P323L)-RdRp.

We consider changes in the equilibrium distributions of the dihedral angles, □, Ψ, and χ_1_. We denote the distribution of a dihedral angle *θ* of the *i-*th residue in wt-RdRp by 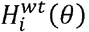,and mt(P323L)-RdRp by 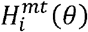. The distributions 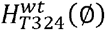 and 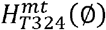, shown in Fig. 3a, are both unimodal but peaks at different locations indicating isomeric transition. This is observed also for 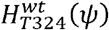 and 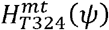 (Fig. 3b) and the side chain dihedrals, 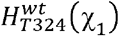 and 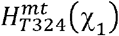 (Fig. 3c). More pronounced changes are observed for F326. 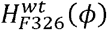 is unimodal whereas 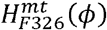 shows bimodal distribution conferring an increase in flexibility at this degree of freedom due to mutation (Fig. 3d). A bimodal distribution is found in 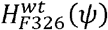, while 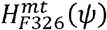is quite flat with multiple peaks (Fig. 3e). 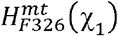is widened more than 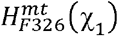 (Fig. 3f). The cases of the dihedral angles of the other residues of the epitope region are shown in SI Figs. S3 and S4.Overall, the distribution of dihedral angles shows a decrease in flexibility due to P323L mutation.

**Figs. 3:**
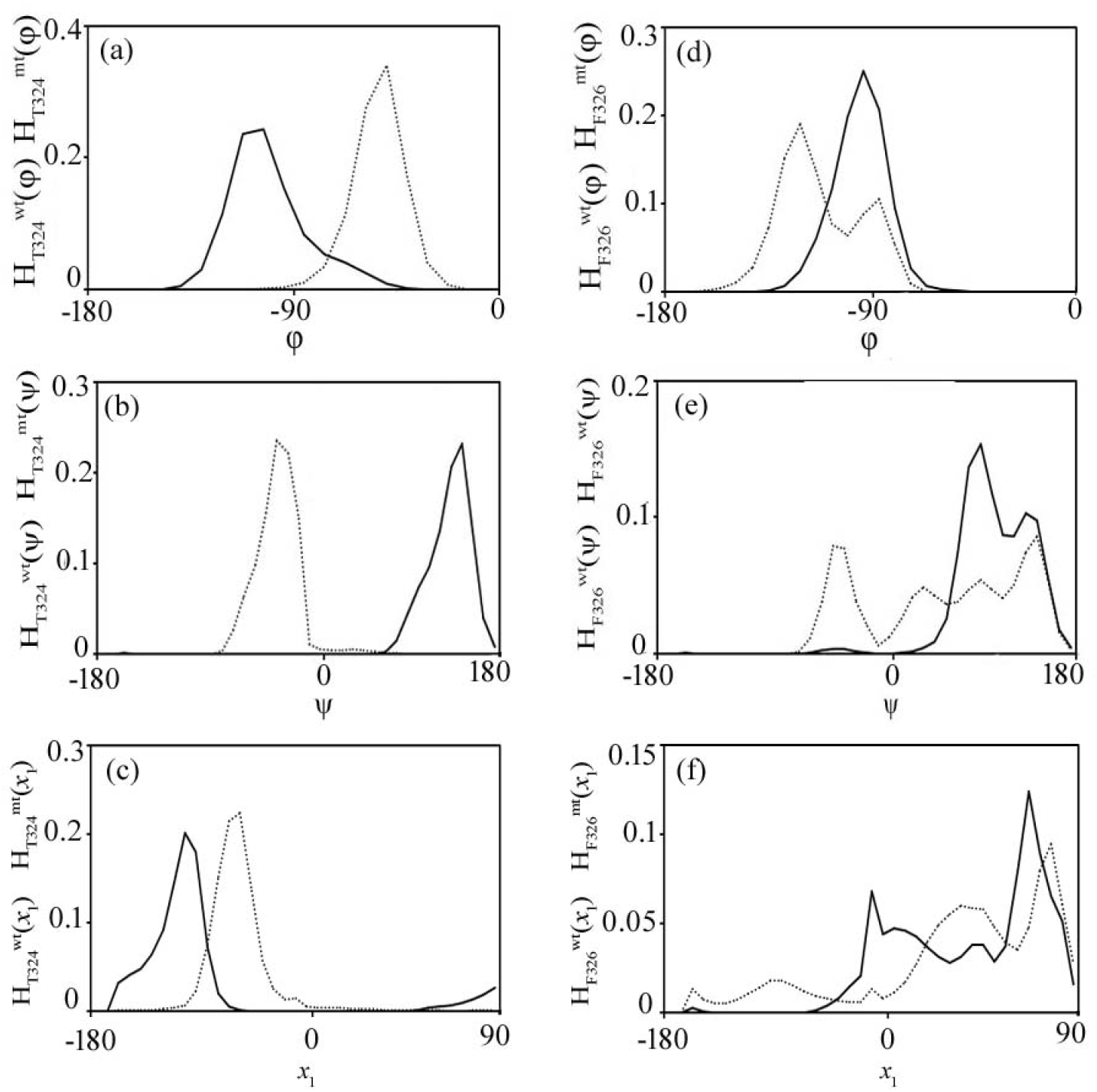
Representative histograms of dihedral angles of wt-RdRp (solid line) and mt(P323L)-RdRp (dotted line) in the epitope. (a) 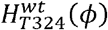 and 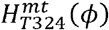. (b) 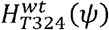 and 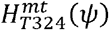. (c) 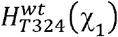 and 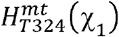. (d) 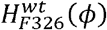 and 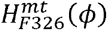. (e) 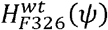 and 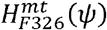. (f) 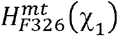 and 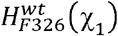.

The changes in the conformational free energy (△G) and entropy (T△S) are computed from comparing the histogram of mt(P323L)-RdRp with respect to wt-RdRp[20]. Careful examination of the epitope region shows that the residues like S325, F326 and G327 undergo enhanced disorder, the maximum being in F326 (Table S1). This is consistent with enhanced flexibilities of the dihedral angles of these residues. On the other hand, there is an increase in order in F321, P322 and T324. The maximum ordering is in T324. The overall change in entropy is 2.53 KJ/mol where the major changes are in the backbone dihedrals. The changes in the stability are marginal for all the cases. Thus, the epitope region undergoes conformational disorder due to P323L mutation of RdRp mainly via enhanced flexibility of the backbone dihedrals.

In case of other point mutations, mt(T324L)-RdRp the epitope conformation is defined by the angle angle*θ* between the vectors joining C_α_atoms of F321 and L324 and G327. H(cos(*θ*)) here is bimodal with peaks at *θ* = 110° and *θ* = 130°. Thus, the epitope prefers more open conformations than in wt-RdRp. In case of mt(F326L)-RdRp, the angle *θ* between the vectors joining C_α_atoms of F321 and P324 and G327, shows a single sharp peak at *θ*= 100°. The mutation here also opens the protein conformation compared to the wild type protein (SI Fig. S5). The conformational free energy and entropy in both cases show destabilization and disorder compared to the wild type. In mt(T342L)-RdRp, the free energy change is 3.30 KJ/mol and the entropy change is noted as 15.86 KJ/mol. The free energy change and entropy change for mt(F326L)-RdRp is 4.71 KJ/mol and 9.24 KJ/mol respectively. Thus, the mutations in the epitope region tend to result in the open conformation of the epitope region and destabilization and disorder compared to the wild type protein.

### 3.2. Changes in epitope-paratope binding interface due to mutation

We carry out docking of the paratope to the epitope with various mutations to ascertain their effects on epitope-paratope binding. The docked complexes are shown in SI Fig. S6. The docking results are shown in Table S2. The binding free energy ΔG and the dissociation constant(K_d_)at 25°C are reported from the docking studies. Both ΔG and K_d_ are less in wt-RdRp-IgG than that in the mutated complexes. Thus, the mutations tend to decrease the binding affinity of IgG. The paratope residues G99 of H1 and W91 of L3 forms van der Waals interactions with the epitope residues T324 and F326 respectively in mt(P323L)-RdRp-IgG.In mt(T324L)-RdRp-IgG complex a single van der Waal interaction is noticed between the epitope residue F326 and L55 of H2 region of paratope. In mt(F326L)-RdRp-IgG complex, we find three hydrogen bonds and one van der Waal interaction in the epitope:paratope interface (Table S3). Thus, the mutations in the paratope region lead to the loss of epitope which impairs the paratope binding. We carry out MD simulations to the microscopic details of impaired paratope binding due to mutation of the paratope residues in the naturally occurring mutation, namely, mt(P323L)-RdRp-IgG and compare the conformational stability with respect to wt-RdRp-IgG. First, we characterize the interface and interactions as revealed by the simulated trajectories. The interfacial residues and remain largely unchanged. We also noted the changes in the nature of interactions between the epitope and paratope interface, listed in Table1. First, we consider the wt-RdRp-IgG complex. A carbon hydrogen bond is formed between P323 (CD:H donor)of wt-RdRp and S32 (O:H acceptor) from L1 region of IgG. The second conventional H-bond interaction is found between L1-S32 (HG: H donor) of paratope and F326 (O: H acceptor) of epitope. The polar epitope residue T324 of wt-RdRp when interacts with charged group H3-D95. The polar epitope residue T324 when interacts with polar paratope residue is responsible for polar interactions (Table 1). The van der Waals interaction energy is about -2.0 KJ/mol. The loss of epitope due to P323L mutation of RdRp disrupts the paratope binding as manifested by a less number of interactions at the mt(P323L)-RdRp-IgG interface (Table1). The paratope residues G99 of H1 and W91 of L3 form van der Waals interactions with the epitope residues T324 and F326 respectively in mt(P323L)-RdRp-IgG. The rest of the CDR from IgG and the associated epitope residues are not found to interact with the interfacial residues of mt(P323L)-RdRp-IgG.

We show the changes in dihedral distributions in dihedral angles in epitope residues in the epitope-paratope complex in Fig. 4. We consider T324 and F326 which participate in binding in both the wild type and mutated complexes. We compare the distributions of the interface residues in the wild type (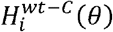,solid line) to those in the mutated 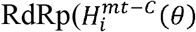,dotted lines).The backbone dihedral *ϕ* in both 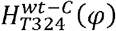 and 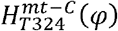 are similar (Fig. 4a). 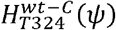 has a tail and is broader than 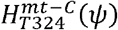 (Fig. 4b). The side chain dihedral (χ_1_) also depicts uniformity in 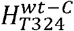 and 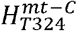 (Fig. 4c). 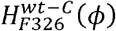 and 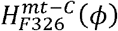 are similar (Fig. 4d). 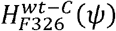 is a unimodal distribution, whereas 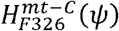 is a multimodal, responsible for the increase in flexibility for this degree of freedom in the epitope:paratope binding interface due to mutation (Fig. 4e). 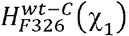 is wider than 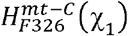 (Fig. 4f). The distributions for additional epitope residues due to mutation are described in SI Figs. S7 and S8.The changes in the dihedral distribution indicate that the epitope residues due to P323L mutation become more flexible.

**Figs. 4:**
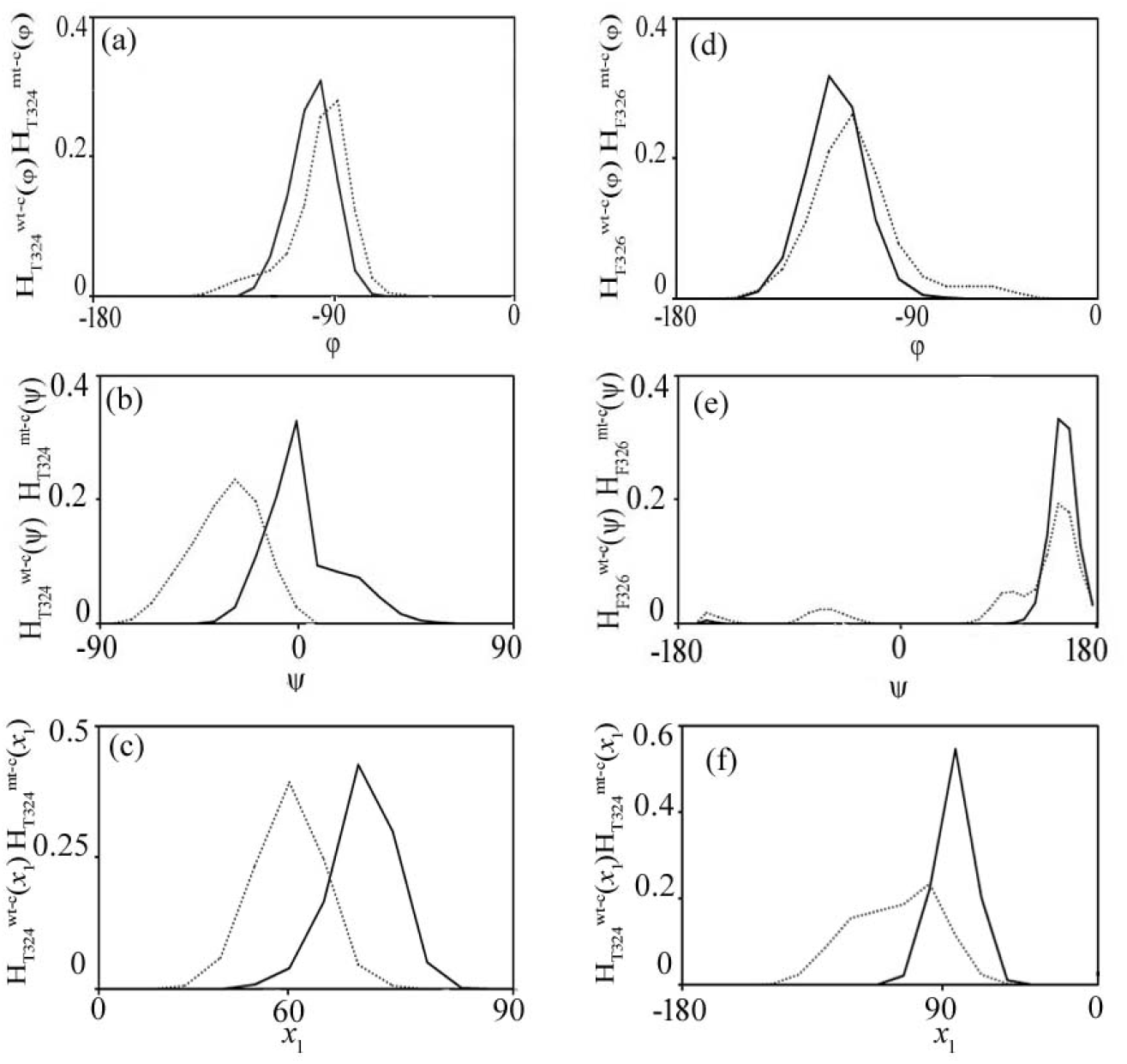
Representative histograms of epitope dihedral angles of the complexes, wt-RdRp-IgG (solid line) and mt(P323L)-RdRp-IgG (dotted line) : (a) 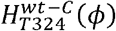 and 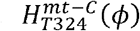. (b) 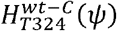 and 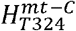. (c) 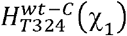 and 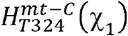 (d) 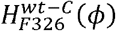 and 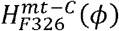. (e) 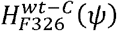 and 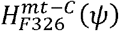. (f) 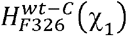 and 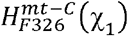.

Some of the changes in distributions of the paratope dihedral angles are shown in Fig. 5. We show distributions:(a) IgG in free state (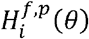,solid line),(b) in bound state of wt-RdRp-IgG (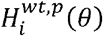, dotted line) and (c) that in mt(P323L)-RdRp-IgG(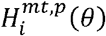, dashed line). 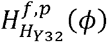 shows a broader distribution than 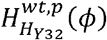 which is sharper, while 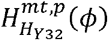 becomes almost as broad as in the free case (Fig. 5a). Thus, the complex in the wild type paratope reduces the flexibility of *ϕ* of the residue but the mutation renders it as flexible as in the free paratope. Similarly, 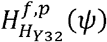 is flatter than 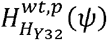, whereas, 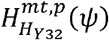 shows a wider distribution similar to 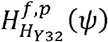 (Fig. 5b). In the case of side-chain dihedral 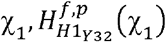, 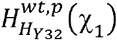 and 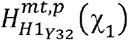 are unimodal with sharp peaks and do not show significant changes (Fig. 5c). From the L chain, 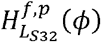 is multimodal but 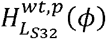 is unimodal, but 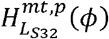 is similar to 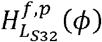 (Fig. 5d). 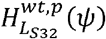 has sharp peak compared to broader peaks in 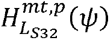 and 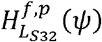 (Fig. 5e). A sharp peak is found in 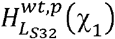, whereas 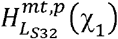 and 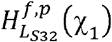 are multimodal in nature (Fig. 5f). The rest of the paratope residues are shown in SI Figs. S9 and S10. Therefore, the flexibility of the free IgG reduces due to epitope: paratope association of the wildtype complex which again increases and become comparable to the free paratope. due to mutation (Fig. 5f). Overall, the changes in dihedral distribution due to the paratope residues indicate that the free unbound paratope loses its flexibility when binding with the epitope region of wt-RdRp.

**Figs. 5:**
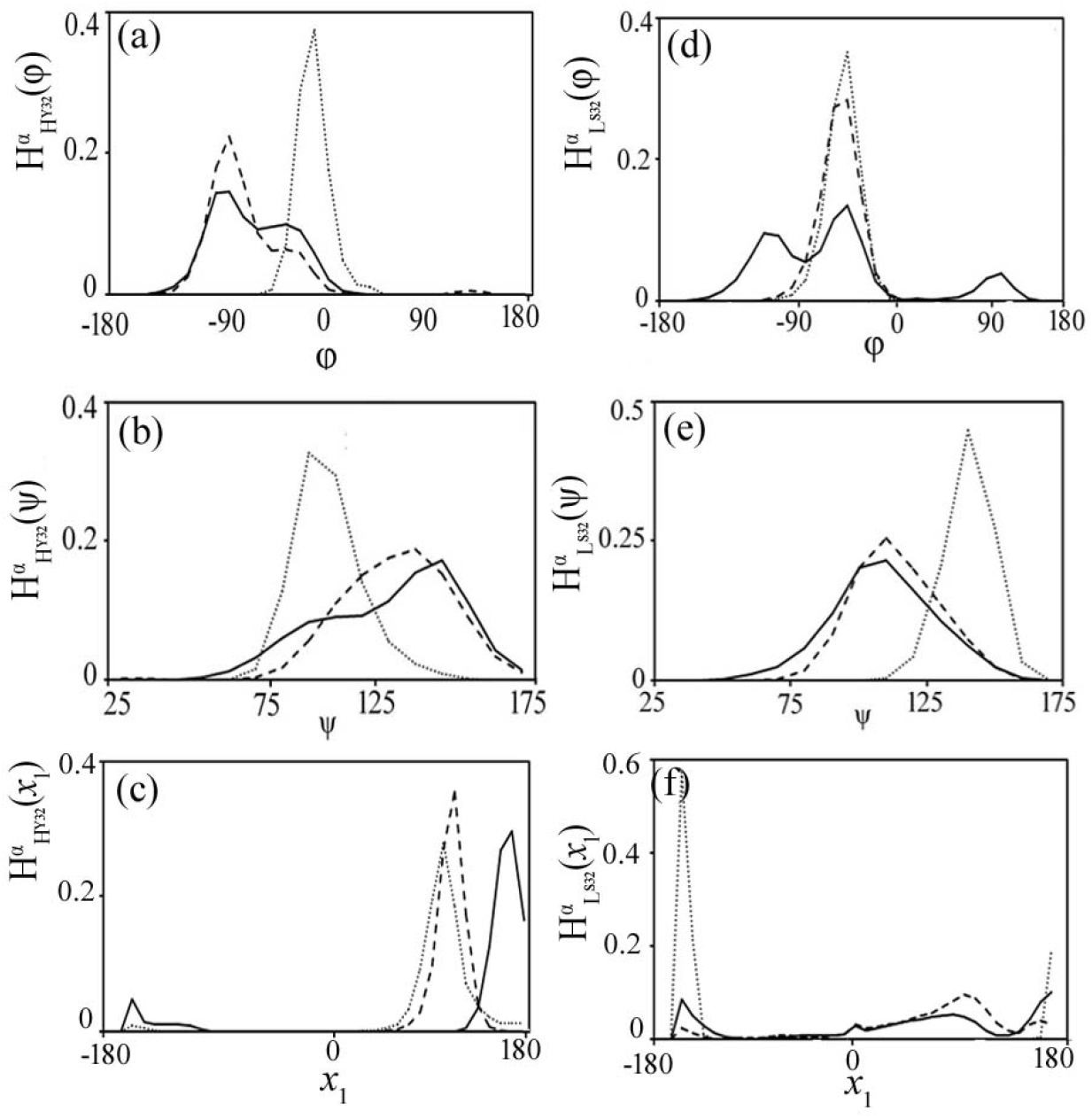
Representative histograms of the paratope dihedral angles IgG in free state (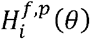,solid line) and in bound state of wt-RdRp-IgG (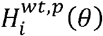, dotted line) and mt(P323L)-RdRp-IgG (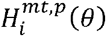, dashed line). (a) *ϕ*, (b) *ψ* and (c) χ_1_ of Y32 of H chain. (d) *ϕ*, (e) *ψ* and (f) χ_1_ of S32 of L chain.

The majority of the residues at the epitope region were found to get ordered and stabilized in wt-RdRp-IgG complex compared to the free state of the epitope in wt-RdRp (Table S4). It is found that F326 is responsible for the largest increase in order. The side chain dihedral (χ_1_) causes a major increase in order. For the change in free energy, the maximum stability is due to T324. The backbone dihedral (*ψ*) contributes most to the increase in stability. Thus, the epitope residues of wt-RdRp are observed to attain stability (−12.93 KJ/mol) and order (−4.04 KJ/mol) due to the paratope binding. On the other hand, binding of mt(P323L)-RdRp with IgG leads to disorder and destabilization in most of the residues compared to those in wt-RdRp (Table S5). The backbone dihedrals are chiefly responsive to the decrease in order. In the case of free energy change due to mt(P323L)-RdRp-IgG complex formation than in the free state of the epitope in wt-RdRp, the residues are marginally destabilized. The backbone dihedrals can be held responsive to the decrease in stability of the region. Thus, the epitope residues in mt(P323L)-RdRp-IgG get destabilized (2.89 KJ/mol) and disordered (4.61 KJ/mol) over wt-RdRp.

We examine the conformational contributions to the free-energy and entropy changes in the paratope residues from the dihedral distributions. First, let us consider wt-RdRp-IgG case (Table S6), in which most of the paratope residues in the complex get ordered and stabilized in the complex compared to their free states. At the paratope region, the backbone dihedral (*ψ*) contributes for the maximum ordering in the complex compared to their free states. Likewise, with respect to free energy change, the maximum stability is imparted by (*ϕ*) dihedral. The backbone dihedrals contribute to the stability of the paratope in wild type complex than the free state. The paratope IgG is found to get ordered (−20.38 KJ/mol) and stabilized (−6.62 KJ/mol) in the wt-RdRp-IgG complex compared to their free states. In the case of mt(P323L)-RdRp-IgG, most of the paratope residues gets disordered and destabilized in the complex than in its free state(Table S7). The mt(P323L)-RdRp-IgG shows the disorder most in all the paratope residues except Y32 and R94 of H chain and S93 of L chain. ExceptingY32 of H chain and W91 and S94 of L chain, all are found to get destabilized due to mt(P323L)-RdRp-IgG complex formation than in its free IgG state. The backbone dihedrals(*ψ*)have a major contribution for the disorder (15.34 KJ/mol) and *ϕ* dihedrals lead to decrease in stability (14.92 KJ/mol) of the mutated complex than in its free state.

## 4. Discussion

The detailed conformational thermodynamics data yield the following microscopic picture of the loss in epitope by a mutation in the epitope region of RdRp. In the mutated protein, the epitope residues are having disordered conformation compared to the wild type protein mostly via the backbone dihedrals, where residues S325 to G327 are primarily affected. The changes in the backbone fluctuations affect the interface with the paratope of IgG. While the stability of the backbone helps to build up a stable interface with the paratope, the enhanced backbone fluctuations in the wild type RdRp disrupt the interactions. The gain in order and stability in the wild type epitope-paratope interface show up in the epitope and paratope residues together both through the backbone and side chain fluctuations. Note that the changes take place all over the epitope-paratope interface, indicating cooperative binding. This is further evident in the wild type case where the epitope-paratope entire region gets disordered and destabilized via both backbone and dihedral fluctuations. Clearly, the mutated RdRp can escape the immune response of the system by altering the backbone fluctuations. Since the conformational disorder can be directly measured by NMR experiments [36,37], the mechanism for immune-escape can be established experimentally.

RdRp is an important therapeutic target to various drug inhibitors [38-44]. Some of the drug molecules are predicted to have binding moiety in a hydrophobic cleft close to the epitope [45]. Thus, it is important to know drug resistance on viral phenotypes due to such mutation. To see the drug resistance at a qualitative level, we dock one drug molecule, Tegobuvir to wt-RdRp and mt(P323L)-RdRp complexes with the grid box center set to (X: 63.99 Å, Y: 91.88 Å, Z: 75.98 Å) and (X: 64.34 Å, Y: 89.74 Å, Z: 74.63 Å) respectively (SI Fig. S11). We find that the binding affinity of Tegobuvir is better for wt-RdRp (Δ G=-9.1Kcal/mol) than that of mt-RdRp (Δ G=-7.9Kcal/mol). Thus, P323L mutation of RdRp alters interactions with Tegobuvir.

To summarize we study here the effect of antibody binding by RdRp due to mutations in the epitope region from MD simulation. We observe that mutations of the epitope residues may cause immune evasion by escaping the antibody-mediated neutralization and producing increased pathogenicity. Albeit the availability of WHO approved emergency vaccines, the impact of SARS-CoV-2 variants of interest and variants of concern on transmissibility, disease severity, risk of reinfection and vaccine performance is a big concern. Thus, such study upon the immune escape variants is a continuous need to understand their epidemiological and clinical impacts. Further vigorous study on the effect of drug molecules on the mutation hot spot of RdRp is essential in future.

## Author contributions

AMG curate analyzed and interpreted the data and wrote the manuscript. SCM helped in writing the codes. SM reviewed and edited the manuscript. JC interpreted the data, reviewed and edited the manuscript.

## Acknowledgements

AMG is thankful to the Technology Research Centre, S.N. Bose National Centre for Basic Sciences, Kolkata for the computational facilities and the Council for Scientific and Industrial Research for financial support through Research Associateship.

